# Bioimaging with fluorescent nucleic-acid aptamers for the specific detection and quantification of *Pseudomonas aeruginosa* alone and in heterogeneous bacterial populations

**DOI:** 10.1101/2025.02.07.636672

**Authors:** Chaimae Mezouarhi, Romain Vauchelles, Basma Abdallah, Janel Régine, Mouna Ouadghiri, Hassan Ait Benhassou, Pierre Fechter, Laurence Choulier

## Abstract

The rising prevalence of bacterial infections, antibiotic resistance, and emerging pathogens underscores the urgent need for innovative diagnostic approaches. Aptamers, short nucleic acid sequences with high specificity and affinity for their targets, are promising candidates for diagnostic applications due to their ability to detect a wide range of pathogens. In this study, we present a fluorescent aptamer-based bioimaging approach for detecting *Pseudomonas aeruginosa*, a multidrug-resistant pathogen of significant clinical concern. Conjugated with fluorescent dye, the detection efficacy of the F23 aptamer was evaluated on 15 different Gram-negative and Gram-positive bacteria, including fixed and lived cells, as well as homogeneous and heterogeneous population. To quantify microscopy images, we developed an automated, open-access identification software using ImageJ. Its high sensitivity provides a robust platform for accurately quantifying bacteria labeled with aptamers and potentially other fluorescent ligands. For instance, it successfully detected 1122 *P. aeruginosa* cells labeled with aptamer F23 out of a total of 1123 *P. aeruginosa* cells in a single image. With almost 200,000 analyzed bacteria and an exceptionally clear signal-to-noise ratio, we demonstrated that the F23 aptamer effectively detects various reference and clinical strains of *Pseudomonas aeruginosa*, while failing to detect Gram-positive *Staphylococcus aureus, Staphylococcus haemolyticus, Staphylococcus epidermidis* and *Corynebacterium striatum*, as well as Gram-negative *Klebsiella pneumonia, Acinetobacter baumannii*, and *Escherichia coli*. The F23 aptamer is therefore a promising tool to distinguish *Pseudomonas aeruginosa* from different strains of the skin microbiota. However, our quantitative method also revealed partial labeling to other bacterial cells, highlighting the issue of refining aptamer selection to improve selectivity.

## 1. INTRODUCTION

The urgent need for improved pathogen surveillance is driven by the growing global population, the increasing prevalence of bacterial infections, the emergence of new pathogens and antibiotic resistance. Accurate and rapid diagnosis methods are therefore becoming increasingly critical (Bergkessel et al., 2023; Rudd et al., 2020). Conventional bacterial diagnosis relies on the culture of bacteria from body samples and the formation of colonies. While highly sensitive, capable of detecting as few as 10^1^ −10^3^ colony forming units per milliliter (CFU/mL), this method is slow, often requiring over 24 hours before staining and observation. Furthermore, it can be imprecise, and error-prone due to sample collection conditions and the specific growth requirements of certain bacteria (Deusenbery et al., 2021). Fluorescence in situ hybridization (FISH) offers an alternative method to identify pathogenic bacteria that bypasses the need for culture. This technique requires fluorescently labeled probes designed to hybridize with complementary sequences of the target organism’s genetic material. When selecting probes for FISH, factors such as specificity, sensitivity and ease of tissue penetration must be considered. Traditional DNA-based FISH often requires permeabilization steps to facilitate probe entry into bacterial cells, which can be time-consuming and may potentially damage samples (Batani et al., 2019; Gu et al., 2022; Moter and Göbel, 2000). Polymerase chain reaction (PCR) is another powerful biotechnological tool for identifying pathogens that are difficult to culture or require long incubation periods (Yamamoto, 2002). Other diagnostic approaches, such as antibody-based biochemical assays and biosensors are also raising interest. These methods often demand specialized equipment, skilled operators, and can be expensive, limiting their accessibility. It is therefore essential to develop alternative diagnosis tools and methods that are simple, cost-effective, rapid, and sensitive (Lin et al., 2023).

Imaging technologies have the potential to significantly enhance bacterial detection by greatly improving both sensitivity and specificity, while enabling more accurate identification and real-time localization of infections. Advanced techniques, such as positron emission tomography (PET) and magnetic resonance imaging (MRI), can be integrated with molecular markers specifically designed to target bacterial components. These molecular probes bind to unique bacterial structures, such as cell walls or metabolic products ensuring precise visualization and differentiation from host cells. The combination of these imaging techniques with bacterial-specific markers allows for more precise monitoring of infection progression and treatment response (Hameed et al., 2018). Moreover, imaging approaches facilitate multiplexed imaging to simultaneously detect multiple bacterial species or biomarkers in a single scan. This capability is particularly beneficial in clinical settings involving complex or polymicrobial infections, where interactions between different bacterial strains and host tissues impact disease progression. Multiplexed imaging provides a comprehensive overview of the infection landscape, delivering critical information for tailored therapeutic interventions and advancing precision medicine in the treatment of bacterial infections (Zhang et al., 2023).

Infections caused by *Pseudomonas aeruginosa* (*P. aeruginosa*) are of particular concern due to several critical factors. This multidrug-resistant opportunistic Gram-negative pathogen exhibits intrinsic resistance to many antibiotics and can rapidly acquire additional resistance through mutations and horizontal gene transfer, further challenging treatment options. Its extensive arsenal of virulence factors, including exotoxins, proteases, and the ability to form biofilms, enhances its capacity to cause severe and persistent infections while evading both antibiotics and the host immune response (Botelho et al., 2019; Qin et al., 2022). Taking advantage of their weakened defenses, *P. aeruginosa* primarily targets immunocompromised individuals, such as cystic fibrosis patients, burn victims, chronic lung diseases sufferers, and hospitalized patients with invasive devices. These infections often lead to severe clinical outcomes, including pneumonia, bloodstream, urinary tract and surgical site complications. Furthermore, the pathogen’s ubiquitous presence in various environments, such as soil, water and moist surfaces in healthcare settings, facilitates the colonization of medical equipment and healthcare personnel, thereby raising the risk of nosocomial infections (Jangra et al., 2022; Tuon et al., 2022). These characteristics highlight the urgent need for stringent infection control measures, and the development of advanced diagnostic approaches to mitigate the impact of *P. aeruginosa* infections.

Nucleic acid aptamers are innovative tools for detection. These short single-stranded DNA or RNA molecules display high affinity and specificity for a wide range of targets, for example small molecules, proteins and whole cells, such as bacteria (Song et al., 2012). This versatility opens up numerous applications in diverse fields including medicine, environmental monitoring, and biotechnology (Zhou and Rossi, 2017). Aptamers also offer several advantages over traditional antibodies, such as ease of chemical synthesis, modification, as well as enhanced stability under diverse conditions, including fluctuations in temperature and pH. Aptamers’ affinity for their targets, generally in the micromolar to low picomolar range, is comparable and sometimes exceeds the performance of monoclonal antibodies (Jenison et al., 1994). In recent years, the use of aptamers as antibacterial agents has gained significant attention due to their potential to fight bacterial infections. They can be engineered to selectively target and bind to bacterial cell surface components, such as adhesins, toxins, or essential enzymes involved in bacterial survival or virulence, offering a potent strategy to address the growing challenge of antibiotic resistance. Aptamers specificity allow them to selectively inhibit the growth or function of pathogenic bacteria without affecting non-targeted cells or tissues (Afrasiabi et al., 2020; Wang et al., 2018). In clinical diagnostics, aptamers can bind to specific bacterial surface proteins or unique bacterial markers, triggering detectable signals that correlate with the presence and concentration of targeted bacteria. This combination of specificity and sensitivity make aptamers particularly valuable for detecting pathogenic bacteria in diverse clinical samples, such as blood, urine, or sputum, thereby enhancing diagnostic precision and enabling timely treatment interventions (Guan and Zhang, 2020). Aptamer-based assays for bacterial toxins detection have gained significant attention. Examples include Aptamer-Linked Immobilized Sorbent Assay (ALISA) kits (Taneja et al., 2020), aptamer-based biosensors (Léguillier et al., 2024), aptamer-based Lateral Flow Assays (LFAs), Enzyme-Linked Aptamer-Sorbent Assays (ELASA) (Rasoulinejad and Gargari, 2016). A key advantage of ELASA systems is regenerability, allowing for repeated use, enhancing the assay’s sustainability and cost-effectiveness. In contrast, antibodies often lack this regenerability, making them less suitable for long-term and cost-effective applications in diagnostics and research settings (Gopinath and Kumar, 2013; Li et al., 2020).

Fluorescence imaging strategies based on nucleic acid aptamers hold particular promise to detect bacteria that are difficult to culture, improving diagnostic accuracy, enabling early diagnosis and facilitating effective monitoring of bacterial infections. Aptamer-based fluorescence imaging strategies has emerged as a powerful tool for the specific detection and visualization of biomolecules, particularly in the context of bacterial infections. These strategies leverage the intrinsic fluorescence of nucleotide bases or modified bases for real-time monitoring of molecular interactions in living cells. Additionally, aptamers can be conjugated with various fluorescent dyes, such as organic dyes, quantum dots, and near-infrared dyes, to enhance imaging performance. A notable advancement in this field is the integration of fluorescence lifetime imaging microscopy (FLIM), which measures the decay time of fluorescence emissions. This approach provides valuable information about the local environment of fluorophores and enables differentiation between bound and unbound states of aptamer-target complexes. FLIM has been successfully employed to detect specific bacterial targets, such as *Staphylococcus aureus* and *Escherichia coli*, using aptamers designed to exhibit altered fluorescence lifetimes upon binding to their targets (Luo et al., 2012; Qazi et al., 2024). Despite their many advantages, including versatility, and the capability for real-time monitoring, challenges remain, such as aptamer stability, non-specific binding, and fluorophore photobleaching. Continued advancements in aptamer design and imaging technologies are essential for expanding their application in biomedical research and diagnostics, particularly in complex biological systems.

In this study, we developed a fluorescence imaging assay for the detection of *P. aeruginosa*, using a single-stranded DNA (ssDNA) aptamer, named F23 (Wang et al., 2011). Additionally, we implemented robust yet user-friendly automated identification technique using the open-access software ImageJ. This approach enables the quantification of *P. aeruginosa* detection from confocal microscopy images of live or fixed stained bacteria, even within heterogeneous bacterial population. Our analytical method is both precise and versatile. This study also raised the question of aptamer’s selectivity against a range of bacterial strains, beyond *Pseudomonas aeruginosa*.

## 2. MATERIALS AND METHODS

### 2.1. Materials

Different bacterial strains were used in this study: *Pseudomonas aeruginosa* ATCC15692 (named *Pa*), *Staphylococcus aureus* HG001, *Bacillus subtilis* ATCC23857, *Escherichia Coli* BW25663, *Pseudomonas aeruginosa* ATCC15692 containing the transposon plasmid pUT18T-miniTn7*gfp* (*Pa-gfp*) encoded gene for Green Fluorescent Protein (GFP) as described (Koch et al., 2001), *Staphylococcus epidermidis* ATCC35983, different clinical isolates of *Pseudomonas aeruginosa* (named *Pa-B5, Pa-B25, Pa-E25, Pa-F1*) (Cramer et al., 2012). *Klebsiella pneumoniae, Acinetobacter baumannii, Staphylococcus haemolyticus, and Corynebacterium striatum* strains were obtained from the collection of Strasbourg University Hospital. *Burkholderia multivorans and Stenotrophomonas maltophilia* were obtained from Anne Doleans-Jordheim collection of Centre International de Recherche en Infectiologie, Université Lyon 1.

SYTO9 green fluorescent nucleic acid stains with an excitation/emission of 485/498 nm, was purchased from ThermoFisher Scientific™ (US). The solution of 4’,6-diamidino-2-phenylindole 2HC1 (DAPI) with an excitation/emission of 359/452 nm and Mueller Hinton II broth (MH) were purchased from Sigma-Aldrich (Hamburg, Germany). The ssDNA F23 aptamer previously described (Wang et al., 2011) was conjugated to cyanine5 (Cy5-F23) at its 5’ end by Eurogentec (Seraing, Belgium).

### 2.2. Bacterial culture preparation

The bacteria strains were inoculated from frozen bacterial stocks. All strains were grown overnight in MH broth without antibiotics, except *Pa-gfp* which was cultivated with 1 μg /ml gentamicin at 37°C under shaking at 220 rpm. The OD at 600 (OD_600_) was adjusted to 0.2 and cells were re-suspended in fresh MH medium. The cultures were then incubated until they reached an OD_600_ of 1. In all experiments, bacteria were diluted when they were in the logarithmic growth phase. The cells were then pelleted by centrifugation at 12,000 rpm for 3 min, to remove the media and, resuspended in fresh MH medium.

### 2.3. Fluorescence-based aptamer-labeling assays on fixed bacteria cells

Bacteria were incubated in 4% paraformaldehyde (PFA) (Sigma Aldrich ®, USA) at room temperature (RT) for 30-60 min. The fixed samples were diluted in 250 mM Glycine for 5-10 min at RT, to ensure efficient quenching. After fixation, cells were washed twice with DPBS 1x (Dulbecco’s Phosphate-Buffered Saline, LONZA), centrifuged at 12,000 rpm for 10 min at RT and resuspended in 500 μL of hybridization buffer (50 mM Tris HCl pH 7.4, 5 mM KCl, 100 mM NaCl, 1mM MgCl_2_ and 0,1% NaN_3_).

All the bacteria used for this study were labeled either with 10 μg/ml of DAPI or SYTO9 dyes in the dark at RT. Excess dye was removed by washing the cells three times with DPBS 1X, and centrifugation at 12,000 rpm for 3 min at RT. The cells were then transferred to microscope slides (Epredia™, Superfrost Plus, Thermo scientific) and dried for 10-30 minutes at 37°C. Figure 1A illustrates the aptamer-based labelling process of fixed bacteria cells.

**Figure 1:**
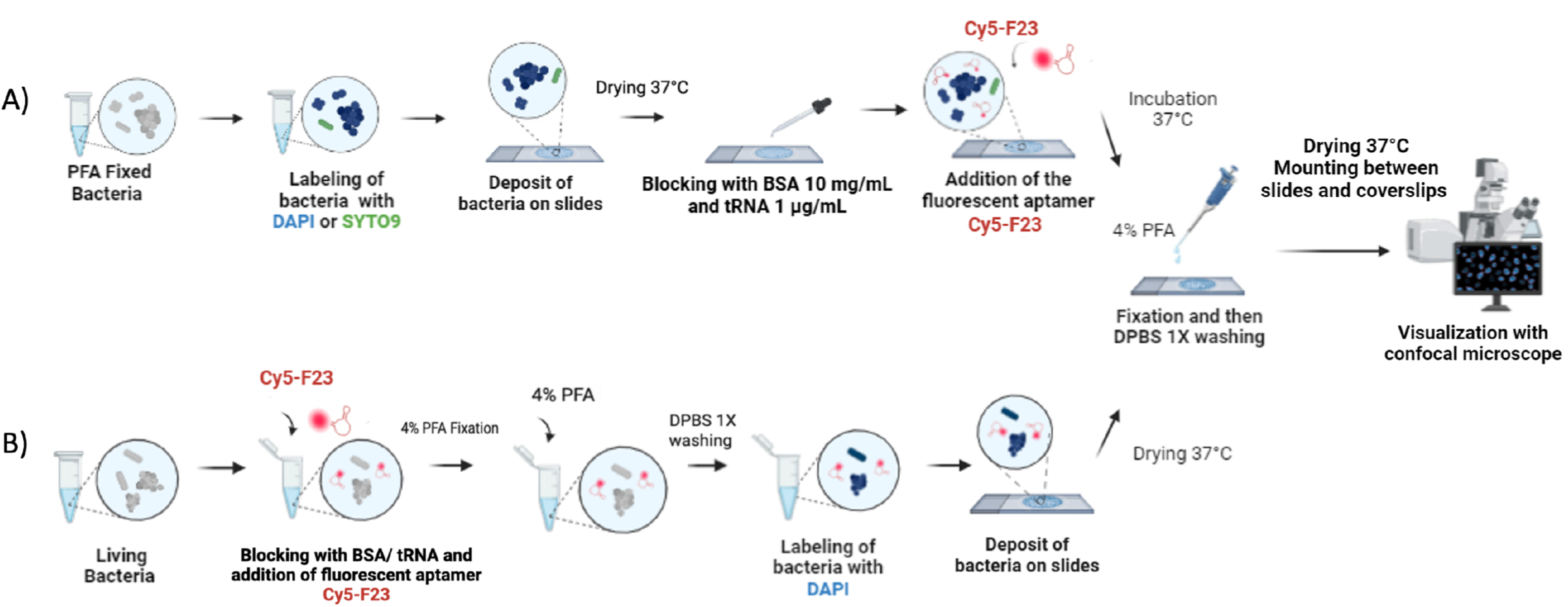
Schematic presentation of the experimental process used for fluorescence-based labeling assays on fixed and live bacteria cells. **A**. Bacteria fixed with 4 % PFA were labeled with DAPI or SYTO9, placed on microscope slides, and dried at 37 °C, treated with hybridization buffer containing BSA and tRNA to block non-specific binding sites and incubated with 1 μM Cy5-F23 aptamer at room temperature (RT). Excess aptamer was removed by washing with DPBS 1x, and the cells were fixed with 4 % PFA, washed, and mounted. **B**. Live bacteria were suspended with hybridization buffer containing BSA and tRNA, incubated with 1 μM of the Cy5-F23 aptamer at RT and then fixed with 4 % PFA before labeling with DAPI. Further steps in the experiment are common to both experimental processes. The cells were then immobilized to microscope slides, before visualization with confocal microscope.

Ten μl of hybridization buffer containing bovine serum albumin (BSA) 10 mg/mL, (Sigma Aldrich) and transfer RNA (tRNA) at 1 μg/mL was added for saturation and incubated for 15 min at RT. The Cy5-F23 aptamer was denatured at 95°C for 5 min, cooled immediately on ice for 5 min, before dilution at 1 μM in hybridization buffer. The Cy5-F23 aptamer was then incubated with fixed cells at RT in a dark room for 60 minutes. After washes with DPBS 1x, 4% PFA was added for 45 min for fixation. After 2 washes with DPBS 1X, and drying, coverslips were mounted using Fluoromount-G™ mounting medium (E140370; Fluoromount-G™, ThermoFisher Scientific™, US).

### 2.4. Fluorescence-based aptamer-labeling assays on living bacterial cells

The protocol for experiments on live bacteria followed the sample procedure as described above with the exception that bacterial strains were not fixed with 4% PFA before aptamer labeling. After culture preparation, bacteria were resuspended in 10 μl of hybridization buffer containing BSA and tRNA, at the above concentrations. Aptamer Cy5-F23 at 1 μM was incubated in Eppendorf tubes with the living cells for 60 min at 37 °C. After incubation, cells were washed with DPBS 1x and centrifuged for 1 min at 12,000 rpm. The bacteria were subsequently fixed with 4% PFA for 45 min and washed. For staining, DAPI was added at 10 μg/mL for 60 min. The samples were then deposited onto microscope slides and dried at 37 °C. Slides were stored in the dark. Figure 1B illustrates this process on living bacteria cells.

### 2.5. Confocal experiments

Images were acquired using a Leica TCS SPE II confocal microscope (Leica Microsystems, Heidelberg, Germany) equipped with HCX PL APO 60x/1.40 oil objective. The confocal microscope was optimally configured for three channels DAPI/SYTO9 or GFP/Cy5 excited with three laser lines with wavelengths of 405 nm/488 nm/635 nm, respectively. Image analysis was performed using macro developed with ImageJ software (Schneider et al., 2012) as described in the result section.

### 2.6. Aptamer stability assay

Aptamer Cy5-F23 at 5 μM in hybridization buffer, MH medium, or MH containing *P. aeruginosa* (OD_600_ of 1) were incubated at 37 °C for 0, 0.5, 2, 5, 10, 24, 48 and 72 hours. Samples were diluted in Blue/Orange Loading Dye (Promega, Madison, WI, USA), denatured at 95°C for 5 min, and loaded on 12% polyacrylamide 8 M urea denaturing gels. Gels were then stained with the Stains all cationic carbocyanine dye (Sigma-Aldrich). Each measurement was performed from three independent experiments. Gel images were taken using Sapphire Biomolecular Imager and analyzed using ImageJ software (v1.54g). All data, analyzed with GraphPad Prism version 5.04, are represented as mean ± SEM.

## 3. RESULTS

### 3.1. Stability Assay

Prior conducting microscopy experiments, we evaluated the stability of the F23 aptamer under different conditions (Figure 2). The F23 aptamer at a concentration of 5 μM was first incubated in the hybridization buffer at 37 °C for 0, 0.5, 2, 5, 10, 24, 48 and 72 h, then loaded on a denaturing 8M urea 12% polyacrylamide gel. The gel displayed a single band corresponding to the full-length aptamer, with no degradation products visible even after 72 hours at 37°C. Quantitative analyses of three independent experiments, confirmed that the F23 aptamer remains highly stable over an extended period of 72h at physiological temperature. While the aptamers were used in hybridization buffer for this study, future applications may involve more complex environments. To explore this, we also tested the stability of the F23 aptamer in Mueller Hinton II medium (MH), commonly used for bacterial culture both alone an in the presence of *P. aeruginosa*. The results showed that the aptamer remains highly stable for 72 hours (Figure 2). This stability at physiological temperature suggests that the Cy5-F23 aptamer could be suitable for extended applications in biological and clinical settings, where consistent performance over time is critical.

**Figure 2:**
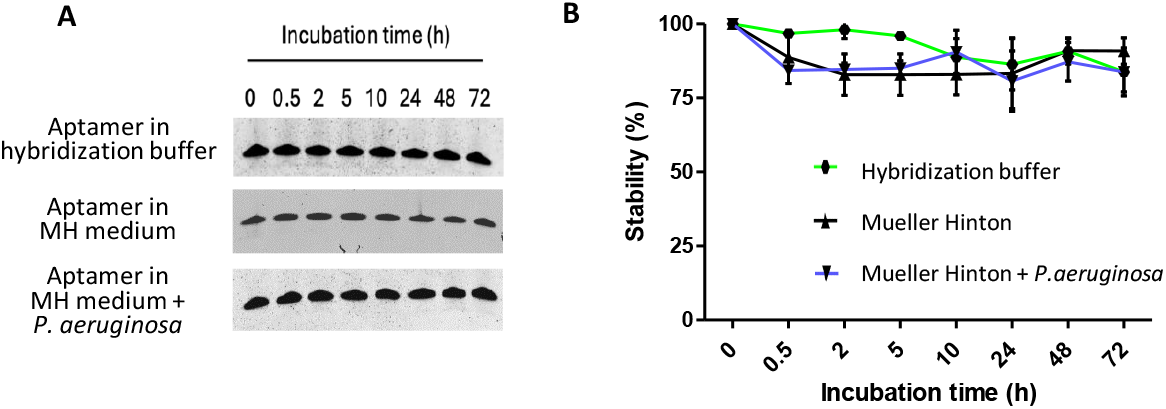
Stability analysis of the Cy5-F23 aptamer. The aptamer Cy5-F23 was incubated at 5 μM for 0, 0.5, 2, 5, 10, 24, 48 and 72 h at 37 °C in hybridization buffer, Mueller Hinton (MH) medium, and MH with *P. aeruginosa*. Samples were loaded on denaturing (8M urea) 12% polyacrylamide gel, revealed using the cationic Stains all solution. **A**. Representative image. One single band indicates that no degradation is detected. **B**. Quantification. Band intensity was quantified using ImageJ at each time point and expressed as a percentage. Data shown are mean ± SD of three independent experiments.

### 3.2. Validation of F23 aptamer binding to *Pseudomonas aeruginosa* strains though confocal imaging

A few studies, such as those by Du et al., Geng et al., and Song et al. (Du et al., 2020; Geng et al., 2021; Song et al., 2017), have depicted aptamer binding to bacterial cells using imaging. Building on this knowledge, our primary objective was to confirm the binding of the ssDNA F23 aptamer (Wang et al., 2011) to *P. aeruginosa* using confocal microscopy. The *P. aeruginosa* strain was fixed using 4% paraformaldehyde, washed, and stained. For staining, we used the two nucleic acid dyes DAPI or SYTO9, both of which are commonly applied to stain live and fixed bacteria. Additionally, we utilized the GFP-labeled strain *Pa* (referred to as *Pa-gfp)* which carries the transposon plasmid pUT18T-miniTn7*gfp* encoding the GFP gene (Koch et al., 2001). Since the different bacterial strains were subsequently combined in cocktails later in the study, different strain were labeled with distinct dyes to enable differentiation. We investigated these staining methods to evaluate whether bacterial labeling influenced aptamer detection and subsequent quantification.

The fixed bacteria were incubated with 1 μM of the F23 aptamer conjugated at its 5’ extremity to the cyanine 5 fluorophore, referred to as Cy5-F23. The bacteria cells were then washed to remove unbound aptamer and imaged using confocal microscopy. The results are presented in Figure 3A-C. Regardless the staining method used (DAPI, SYTO9 or GFP), the images demonstrated strong and overlapping fluorescence signals from the aptamer and the DAPI/SYTO9/GFP labeled *P. aeruginosa*, indicating highly effective binding of the F23 aptamer. In the merged images, the colocalization of the F23 aptamer and *P. aeruginosa* cells was evident. Yellow signals were observed when green fluorescence (from SYTO9 or GFP-labeled *P. aeruginosa*) overlapped with the red fluorescence of the Cy5-F23 aptamer (Figure 3A and 3C). Similarly, violet signals resulted from the superposition of blue fluorescence (DAPI-stained *P. aeruginosa*) with the red Cy5-F23 aptamer fluorescence (Figure 3B). A few red-only cells in the merged images corresponded to *P. aeruginosa* labeled by the Cy5-F23 aptamer but either unlabeled or weakly labeled with SYTO9, highlighting the aptamer’s robust binding and labeling capabilities even in the absence of strong SYTO9 staining. To assess selectivity, the binding of the Cy5-F23 aptamer to *Staphylococcus aureus* (*S. aureus*) was also evaluated using the same methodology. As shown in Figure 3D, no fluorescence signal was detected in the Cy5 channel, confirming that the Cy5-F23 aptamer did not bind to *S. aureus*. Due to the highly distinct fluorescence patterns observed with *P. aeruginosa* and *S. aureus* bacteria, we selected the F23 aptamer for further investigation of its specificity against additional bacterial strains and for quantification. For this purpose, we developed the macro described below.

**Figure 3:**
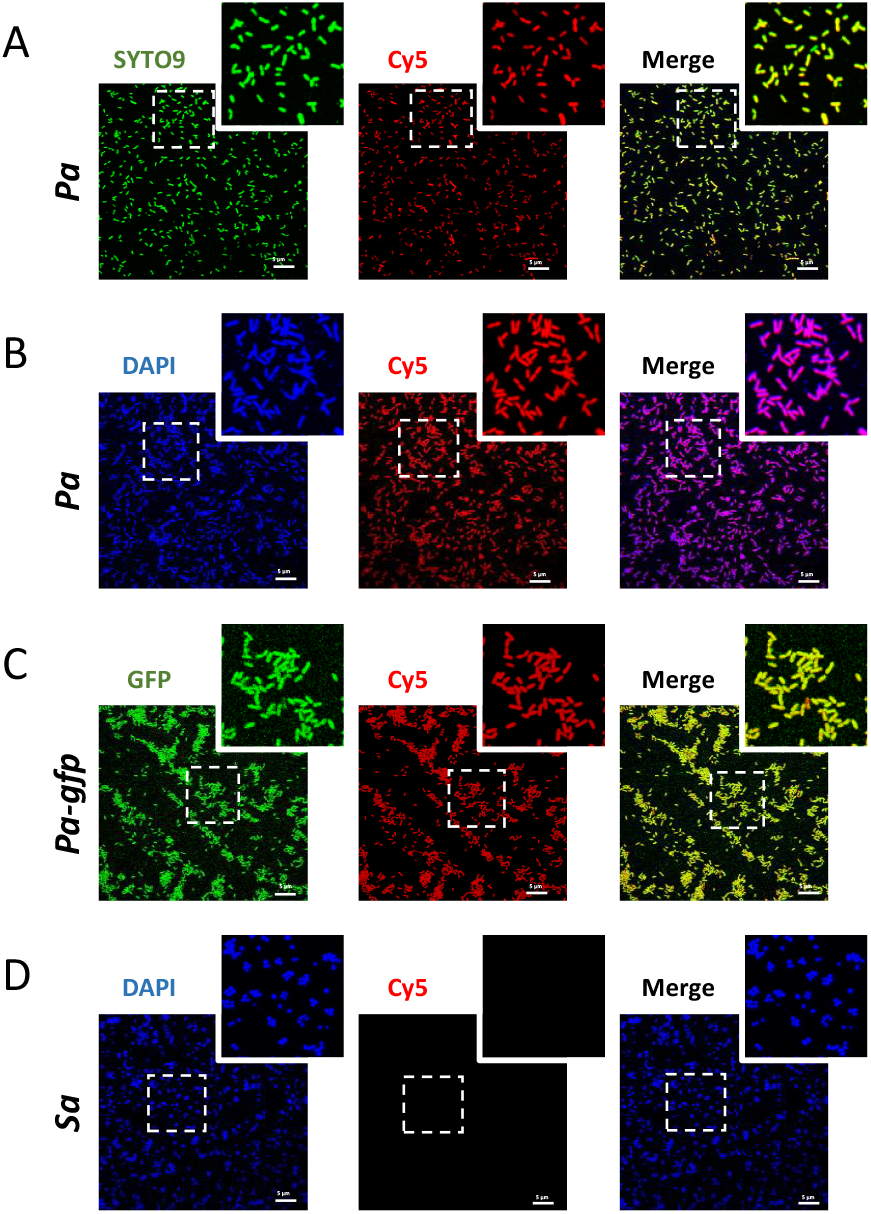
Representative confocal images of F23 aptamer on the two bacteria strains *P. aeruginosa* and *S. aureus*. One μM of the F23 aptamer conjugated to the cyanine-5 fluorophore (Cy5-F23) was incubated for 60 min with fixed strains of *Pseudomonas aeruginosa*, referred as *Pa* (**A-C**) and *Staphylococcus aureus*, referred as *Sa* (**D**). *Pa* was stained with **A**. SYTO9 (λex = 498 nm, represented in green), **B**. DAPI (λex = 412 nm, represented in blue). In **C**. the GFP-tagged *Pa* strain, *Pa-gfp*, was used (λex = 498 nm, represented in green). In **A-C**, the positive binding of the Cy5-F23 aptamer on *Pa* is represented in red (Cy5, λex = 644 nm). On the merged channels, the yellow (**A** and **C**) or violet (**B**) colors show the superposition of the fluorescence emitted by the dyes or GFP with that in red emitted by the F23 aptamer. The rare red bacteria which subsist on merged images represents *Pa* labeled by the F23 aptamer but either not marked or only weakly marked by dyes or GFP. **D**. *Sa* stained with DAPI (represented in blue). No fluorescence is emitted in the Cy5 channel because the Cy5-F23 aptamer does not bind to *Sa*. **A-D**. Images were acquired with a Leica TCS-SPE confocal microscope. Scale bar: 5 μm. Magnified images are from the inserts.

### 3.3. Macro development for the quantification of aptamers labeling bacteria using ImageJ, with code availability

A set of imageJ macros that requires ImageJ v1.54g or higher was developed to analyze each image and quantify aptamer binding on stained bacteria. The workflow for using the macro is illustrated in Figure 4. Source code and documentation for the installation and use of macros are available at in supplementary data.

**Figure 4.**
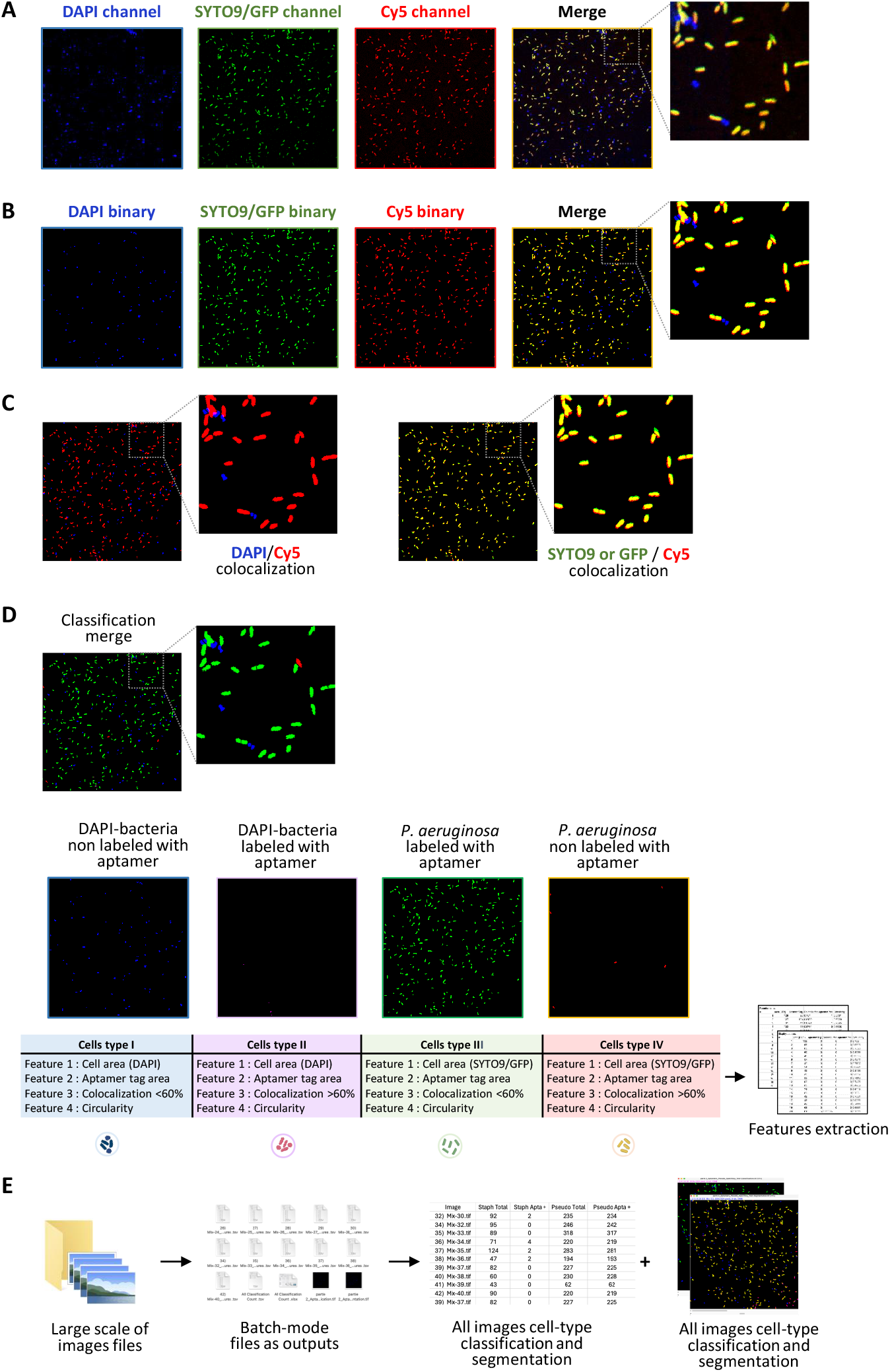
Schematic representation of the custom-developed macros used for the quantifying confocal microscopy images obtained from the aptamer-based labeling of bacterial cells. **A**. Images from confocal microscope are acquired and then processed for analysis using the open-source ImageJ. **B**. Image segmentation: segmentation is performed using the adaptive thresholding method for DAPI and SYTO9/GFP channel and classic thresholding for Cy5 channel. Identification numbers are assigned to individual bacteria, regions of interest (ROIs) are extracted and saved in the ROI manager. **C**. Colocalization of resulting binary images: surface overlap is measured between bacteria tag (DAPI or SYTO9/GFP) and aptamer label (Cy5) for each bacteria. **D**. Bacterial cell classification: bacteria are classified into the following classes: I. DAPI-bacteria non labelled with aptamer, II. DAPI-bacteria labelled with aptamer, III. *P. aeruginosa* labelled with aptamer, IV. *P. aeruginosa* non labelled with aptamer. Feature extraction is performed and quantification results in .tsv file format of classification cells belonging to each class is given. **E**. Batch mode. The analysis can also be run in batch mode to process large image sets. All segmented and classified cell-type images are included in the output.

All macros share the same segmentation technique inspired by the adaptative thresholding method (Abdolhoseini et al., 2019). Images were first denoised using a gaussian filter, followed by background subtraction. Local maximas were then defined and used as seeds for growing regions. Each resulting area was binarized using its own threshold defined by the Otsu algorithm. A watershed algorithm was applied to separate touching bacteria. A size filter was used to exclude large regions containing several bacteria. The densely confluent parts of the bacteria population were excluded from this analysis.

A first classification macro (*Aptamers_Pseudo_Specificity_test*.*ijm)* was developed to assess the selectivity of aptamer for *Pseudomonas* bacteria. This macro was applied to *Pseudomonas* (SYTO9/GFP channel) and other selected strains (DAPI channel) both individually and in mixtures, to evaluate the aptamer’s ability to identify bacteria (Cy5 channel). The DAPI and GFP channels were both segmented using an adaptative threshold method. An aptamer positive mask was then created using a simple Otsu threshold. Bacteria were considered positive for aptamer labelling if 60% or more of their area overlap this mask. The 60% threshold was set considering the bacteria drift due to convection in the media between the acquisition of different channels.

The second macro (*Aptamers_Pseudo_Classifier*.*ijm*) was designed to spot *P. aeruginosa* in a mixed bacterial population, in which the whole heterogeneous bacterial mixture was labelled with DAPI and Cy5-aptamer. Images were segmented and classified using the same method as described for the first macro. Identification numbers were assigned to individual bacteria, regions of interest (ROIs), were extracted and saved separately in ROI manager, and classification categories were quantified.

Binding efficiency of the aptamer to bacterial cells was then analyzed by comparing labeled and unlabeled groups, with statistical analysis used to determine significance. For large datasets, batch processing was implemented to automate the analysis. Results, including segmented images, ROIs, and classification data, were saved, and a report was generated, documenting all parameters used for reproducibility, alongside the images.

### 3.4. Bioimaging detection and quantification of different fixed bacteria strains

The Cy5-F23 aptamer was evaluated for its binding capacity to bacterial strains beyond *P. aeruginosa* (*Pa*) and *S. aureus* (*Sa*), in order to assess its broader potential for bacteria detection. In detail, the selectivity of Cy5-F23 was tested on the following strains: (i) the model Gram-positive and Gram-negative strains *Bacillus subtilis* (*Bs*) and *Escherichia coli* (*Ec*), (ii) some of the major bacteria from the skin microbiota *Staphylococcus epidermis* (*Se*), *Staphylococcus haemolyticus* (*Sh*), *Corynebacterium striatum* (*Cs*), (iii) different clinical isolates of the Gram-negative bacteria strains *Pseudomonas aeruginosa* referred as *Pa-B5, Pa-B25, Pa*-*E25, Pa*-*F1*,, and (iv) different strains involved in infections of Cystic Fibrosis patients like *Burkholderia multivorans* (*Bm*), *Stenotrophomonas maltophilia* (*Sm*), *Klebsiella pneumonia* (*Kp*), *Acinetobacter baumannii* (*Ab*).

All *Pseudomonas* strains were either stained with SYTO9 or were GFP-labeled, while the remaining bacterial strains were stained with DAPI. The binding capacity of the Cy5-F23 aptamer was examined on these strains. Figure 5 shows representative single confocal images for each tested bacterial strain, alongside corresponding quantitative binding data analyzed using the custom macro described in the previous chapter. Figure 6 summarizes the overall quantification, based on the analysis of all analyzed images, corresponding to 3,000-32,000 bacteria quantified / strain. The results, as shown in Figures 5 and 6, are organized into three groups based on the F23 aptamer bacterial-binding efficiency: A. low binding, B. high binding, and C. moderate binding. This comprehensive analysis highlights the variability of the Cy5-F23 aptamer’s binding across different bacterial species, providing valuable insights into its potential application for bacterial detection.

**Figure 5:**
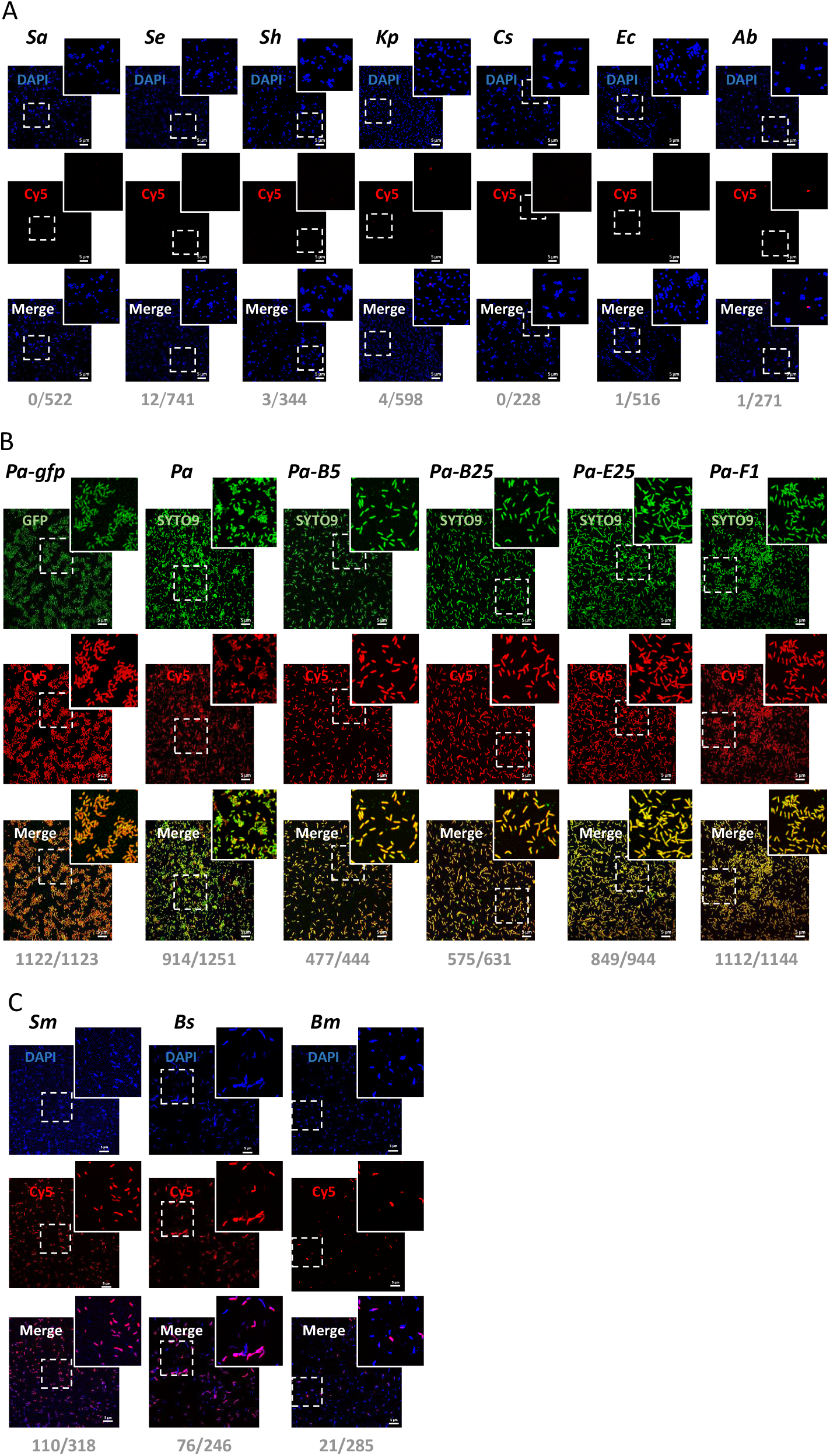
F23 aptamer binding assay on different bacterial strains. Confocal microscopy images and quantification on **A**. bacteria which bind weakly to the Cy5-F23 aptamer: *Staphylococcus aureus* (*Sa*), *Staphylococcus epidermidis* (*Se*), *Staphylococcus haemolyticus (Sh), Klebsiella pneumoniae* (*Kp*), *Corynebacterium striatum (Cs), Escherichia coli* (*Ec*), *Acinetobacter baumannii* (*Ab*), **B**. bacteria which bind strongly the Cy5-F23 aptamer: *Pseudomonas aeruginosa (Pa)*, GFP-tagged *Pa* (*Pa-gfp*) and different *Pseudomonas aeruginosa* clinical isolates referred as (*Pa-B5, Pa-B25, Pa-E25, Pa-F1*), **C**. bacteria which bind partially to the Cy5-F23 aptamer: *Bacillus subtilis* (*Bs*), *Stenotrophomonas maltophilia* (*Sm*), *Burkholderia multivorans* (*Bm*). **A-C**. Briefly, 1 μM of the Cy5-F23 aptamer was incubated for 60 min with fixed bacterial strains. Images were acquired with a Leica TCS-SPE confocal microscope. All *Pseudomonas* strains were stained in green (SYTO9/GFP, λex = 498 nm) and all the other strains were stained in blue (DAPI, λex = 419 nm). A positive binding of the aptamer is shown in red (Cy5, λex = 644 nm). The numbers indicate the ratio of the number of aptamer-labeled bacteria for each strain to the total number of bacteria analyzed in the image (e.g., 0/522 for *S. aureus*). Scale bar: 5 μm. Magnified images are from the inserts.

**Figure 6.**
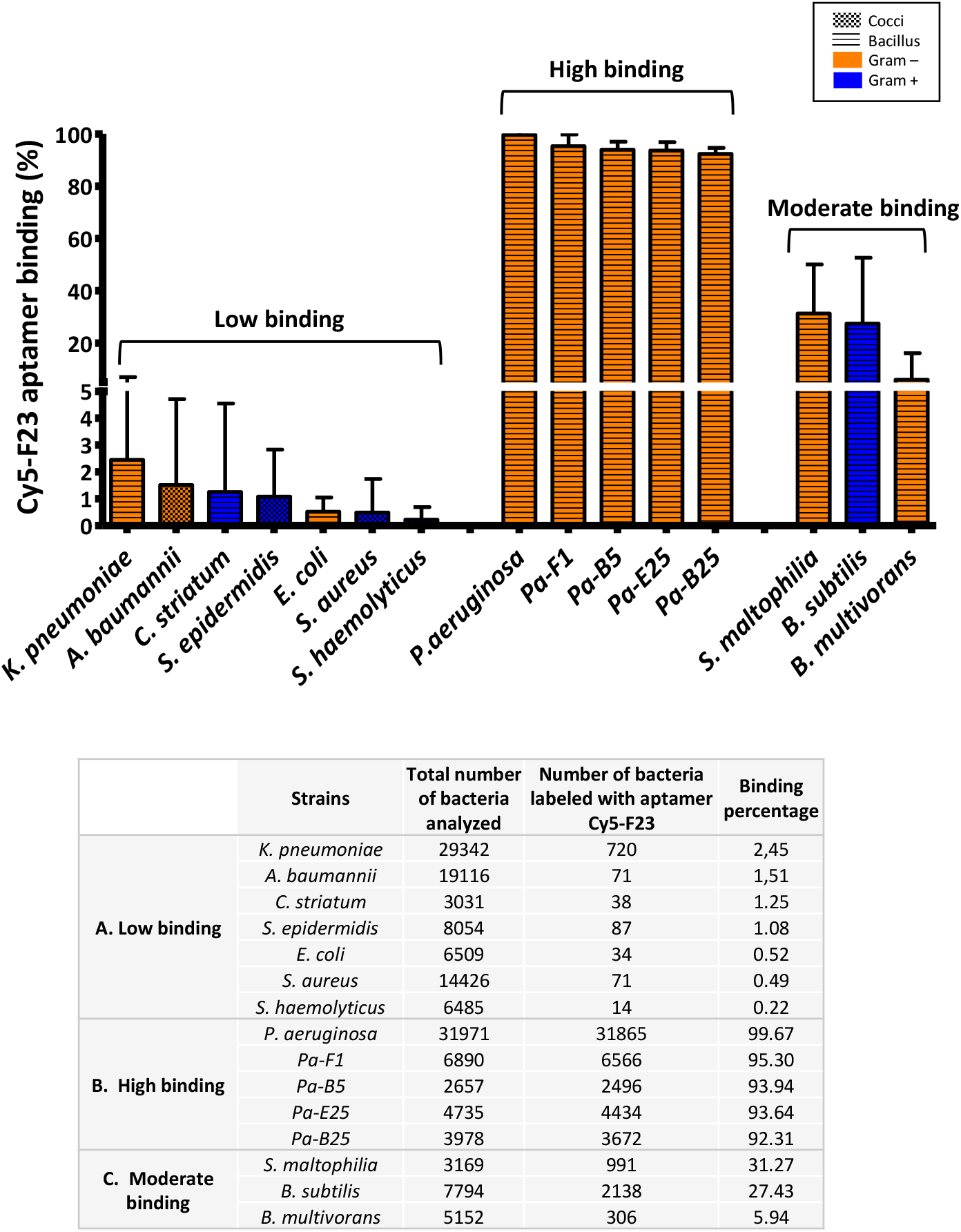
Global quantification of the Cy5-F23 aptamer binding on different bacterial strains. Binding percentages were calculated from the number of cells labeled with the Cy5-F23 aptamer relative to the total number of cells analyzed. Bacterial strains are classified in 3 groups as defined in the text: A. Low binding, B. High binding and C. Moderate binding. Bacterial strains are also distinguished by two colors: blue for Gram-negative strains and orange for Gram-positive strains, and bacterial shapes are distinguished by two patterns: gingham for rod-shaped and horizontal stripes for round bacteria. The table summarizes the number of cells analyzed and the corresponding aptamer-binding percentages for each strain. *P. aeruginosa* (*Pa*) exhibited the highest binding efficiency (99.67%), while *S. haemolyticus* (*Sh*) had the lowest (0.22%).

Bacteria with low binding (below 5%) to the F23 aptamer (Figures 5A and 6A) include *S. aureus* (as already shown in Figures 3), *S. epidermidi*s, *S. haemolyticus, E. coli, C. striatum, K. pneumoniae*, and *A. baumannii*. The Cy5-F23 aptamer does not bind to any tested *Staphylococcus* strain. For *S. aureus*, DAPI staining revealed numerous blue-fluorescent bacteria, confirming the presence of bacterial DNA. However, the Cy5 and merged fluorescence channels showed no evidence of aptamer labeling, with only DAPI-induced blue fluorescence observed. Quantitatively, none of the 522 analyzed *S. aureus* cells were labeled by the aptamer (0/522). Similar results were observed for the other strains: *S. epidermidis* (12/741), *S. hemolyticus* (3/344), *K. pneumoniae* (4/598), *C. striatum* (0/228), *E. coli* (1/516), and *A. baumannii* (1/271). The global quantitative analysis conducted on a larger set of images (Figure 6A) corroborates this observation showing negligible binding of the Cy5-F23 aptamer to these strains: 2.45% for *K. pneumoniae*, 1.51% for *A. baumannii*, 1.25% for *C. striatum*, 1.08% for *S. epidermidis*, 0.52% for *E. coli*, 0.49 % for *S. aureus*, and 0.22% for *S. haemolyticus*.

All *P. aeruginosa* strains exhibited green fluorescence with consistent fluorescence patterns observed across different genotypes (Figure 5B). The merged images displayed yellow-orange fluorescence, resulting from the overlap of the green signal (*Pa*) and the red signal (Cy5-F23 aptamer), confirming the colocalization of the aptamer with *P. aeruginosa* cells. In some regions of the merged images, red fluorescence was observed, corresponding to *P. aeruginosa* cells bound by the Cy5-F23 aptamer but either unlabeled or only weakly labeled with SYTO9. This finding highlights the aptamer’s ability to label *P. aeruginosa* even when SYTO9 staining is low or absent, as previously noted. The quantitative analysis of all confocal images (Figure 6B) revealed that out of a total of 31,971 *P. aeruginosa* cells examined, 31,865 were labeled by the F23 aptamer, representing a very high binding rate of 99.67%. For the clinical strains, the binding rate is 92 to 95 %. These results underscore the high specificity of the Cy5-F23 aptamer for detecting *P. aeruginosa*.

Among the group showing low to moderate binding of the F23 aptamer (Figure 5C) with efficiencies comprised between 5% and 35%, are *B. subtilis* (76/246), *S. maltophilia* (*Sm*, 110/318), and *B. multivorans* (21/285). The global results obtained from analyzing all images (Figure 6C) support these observations, with aptamer-labeling rates of 31.27 % for *S. maltophilia*, 27.43 % for *B. subtilis*, and 5.94 % for *B. multivorans*.

These results highlight the variability of the F23 aptamer’s binding between different bacterial strains. The observed discrepancies may result from differences in surface structures or molecular targets among the strains, which could influence the aptamer’s recognition or accessibility. Gaining a deeper understanding of structural and molecular variations is likely crucial for optimizing the use of aptamers in strain-specific detection or therapeutic applications.

### 3.5. Bioimaging detection and quantification of heterogeneous bacteria populations in mixed samples

Having demonstrated that the F23 aptamer can accurately distinguish between *P. aeruginosa* and *S. aureus* in homogeneous samples, we next used the *Aptamers_Pseudo_Classifier*.*ijm* macro to assess its ability to identify its bacterial target within a heterogeneous bacterial population.

In these experiments, *P. aeruginosa* were stained with SYTO9, while other bacterial cells were stained with DAPI. Following staining, bacteria were washed to remove excess dyes, mixed, placed on the microscope slide, incubated for 1 hour with the Cy5-F23 aptamer at room temperature and then mounted for visualization under the microscope.

In a heterogeneous community of *P. aeruginosa* and *S. aureus* (Figure 7A), the F23 aptamer exhibited selective binding to *P. aeruginosa*. Red fluorescence resulting from the binding of the Cy5-F23 aptamer was observed exclusively for *P. aeruginosa* cells, which appeared yellow-orange in the merged image. Quantitative analysis revealed strong binding with 255 out of 278 *P. aeruginosa* cells labeled with aptamers. In contrast, *S. aureus* displayed only blue fluorescence from DAPI staining with no red signal from the Cy5-F23 aptamer (0/122). Across all analyzed images (Table in Figure 7), only 6 out of 1,258 *Sa* cells (0.46%) showed aptamer labeling, compared to 3,300 out of 3,531 *P. aeruginosa* cells (93.46%). These results confirm the high specificity of the F23 aptamer for *P. aeruginosa*, and its lack of labeling of *S. aureus*, even within heterogenous bacterial populations.

**Figure 7.**
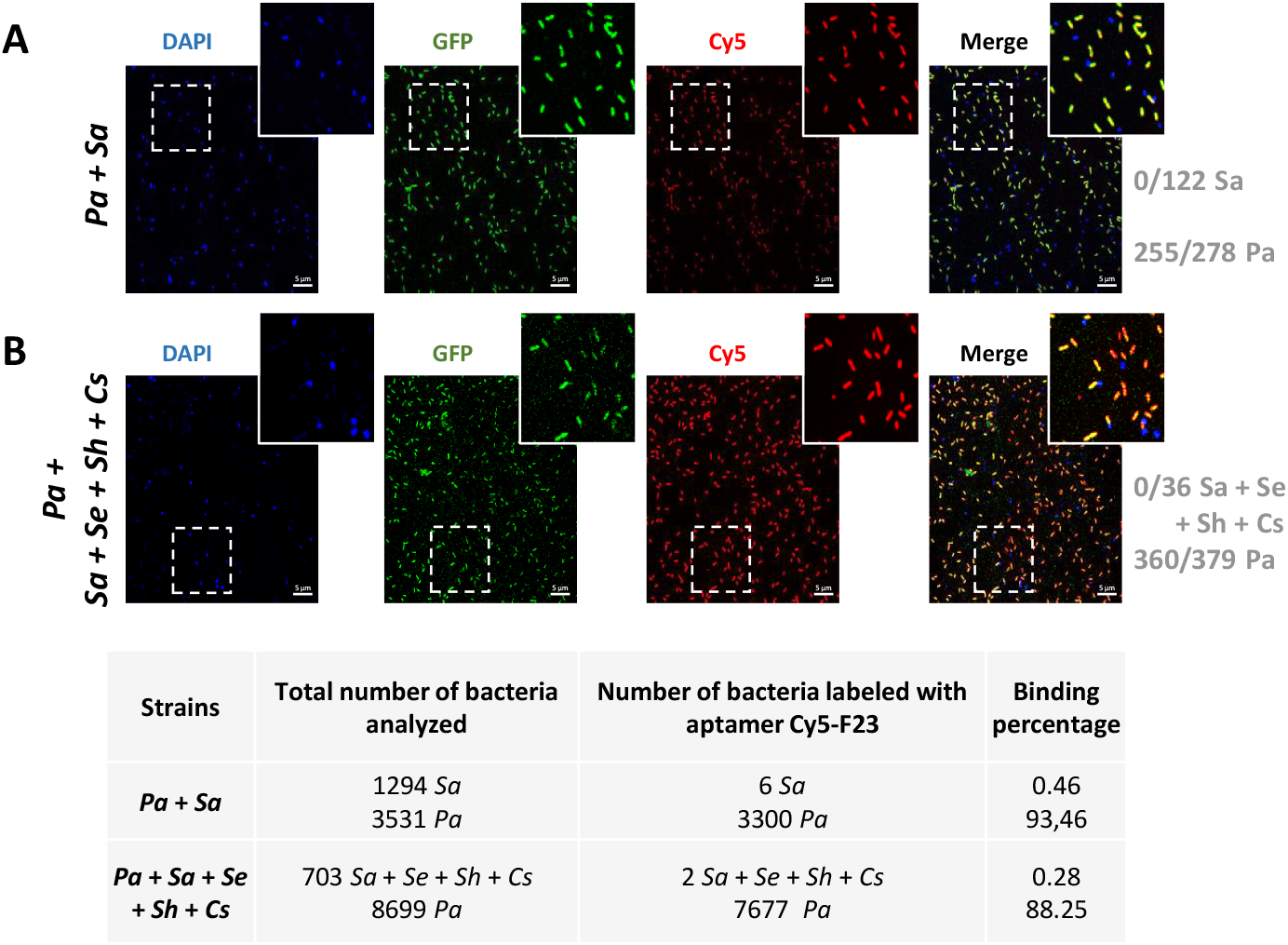
Confocal microscopy imaging of the F23 aptamer on heterogeneous populations of fixed bacteria strains. *P. aeruginosa* (*Pa*) was labelled with SYTO9 (represented in green), and all the other bacteria used for this experiment were labelled with DAPI (represented in blue). One μM of the Cy5-F23 aptamer was incubated for 60 min with a cocktail of different strains: **A**. *P. aeruginosa* (*Pa*) and *S. aureus* (*Sa*). **B**. *P. aeruginosa* (*Pa*), *S. aureus* (*Sa*), *S. epidermidis* (Se), *S. haemolitycus* (*Sh*), *C. striatum* (*Cs*). A positive binding with the aptamer in the merged images is shown in yellow-orange color. The F23 aptamer only binds to *P. aeruginosa* in these mixture of bacteria. Scale bar: 5 μm. Magnified images are from the inserts. The table lists the total number of cells analyzed and the corresponding aptamer-binding percentages for each strain.

The selectivity of the F23 aptamer was further validated in a more complex mixture, where *P. aeruginosa* cells were stained with SYTO9, while *S. aureus, S. epidermidis, S. haemolyticus* and *C. striatum* were stained with DAPI (Figure 7B). The F23 aptamer exclusively bound to *P. aeruginosa*, with no Cy5 labeling observed for the other bacterial strains, which remain distinctly blue. The yellow-orange fluorescence in the merged image confirms the specific binding of the F23 aptamer to *P. aeruginosa*. These results are quantitatively supported by binding rates of blue stained bacteria *S. aureus, S. epidermidis, S. haemolyticus, C. striatum*, with 0 out of 36 cells labeled with the Cy5-F23 aptamer. In contrast, the aptamer effectively targeted nearly all *P. aeruginosa* cells, with a binding rate of 360 out of 379 cells. The macro analysis further underscores the potential of the Cy5-F23 aptamer for detecting over 88 % of *P. aeruginosa* in heterogeneous bacterial populations, demonstrating its high specificity and suitability for complex sample environments.

### 3.6. Bioimaging of F23 Aptamer on living bacterial strains

Experiments were conducted on live DAPI-labeled *P. aeruginosa* and *S. aureus* strains to evaluate whether the F23 aptamer maintained its binding ability with both PFA-fixed and live bacteria. For live cell experiments, *P. aeruginosa* and *S. aureus* cells were incubated at RT for 1 hour with the Cy5-F23 aptamer at 1 μM, before fixation and DAPI-labeling.

The image analysis (Figure 8A-B) confirms the specificity of the F23 aptamer for *P. aeruginosa*. The aptamer labeled only the DAPI-stained *P. aeruginosa*, with 305 out of 383 cells showing positive binding. In contrast, no aptamer-binding was detected on live *S. aureus* (0/277). Quantitative analysis of 4,204 *P. aeruginosa* and 6,767 *S. aureus*, further support these findings: 73.79% of live *P. aeruginosa* cells (3102/4204) were labelled by the Cy5-F23 aptamer, compared to only 0.06% of live *S*.*aureus* (4/6,767).

**Figure 8.**
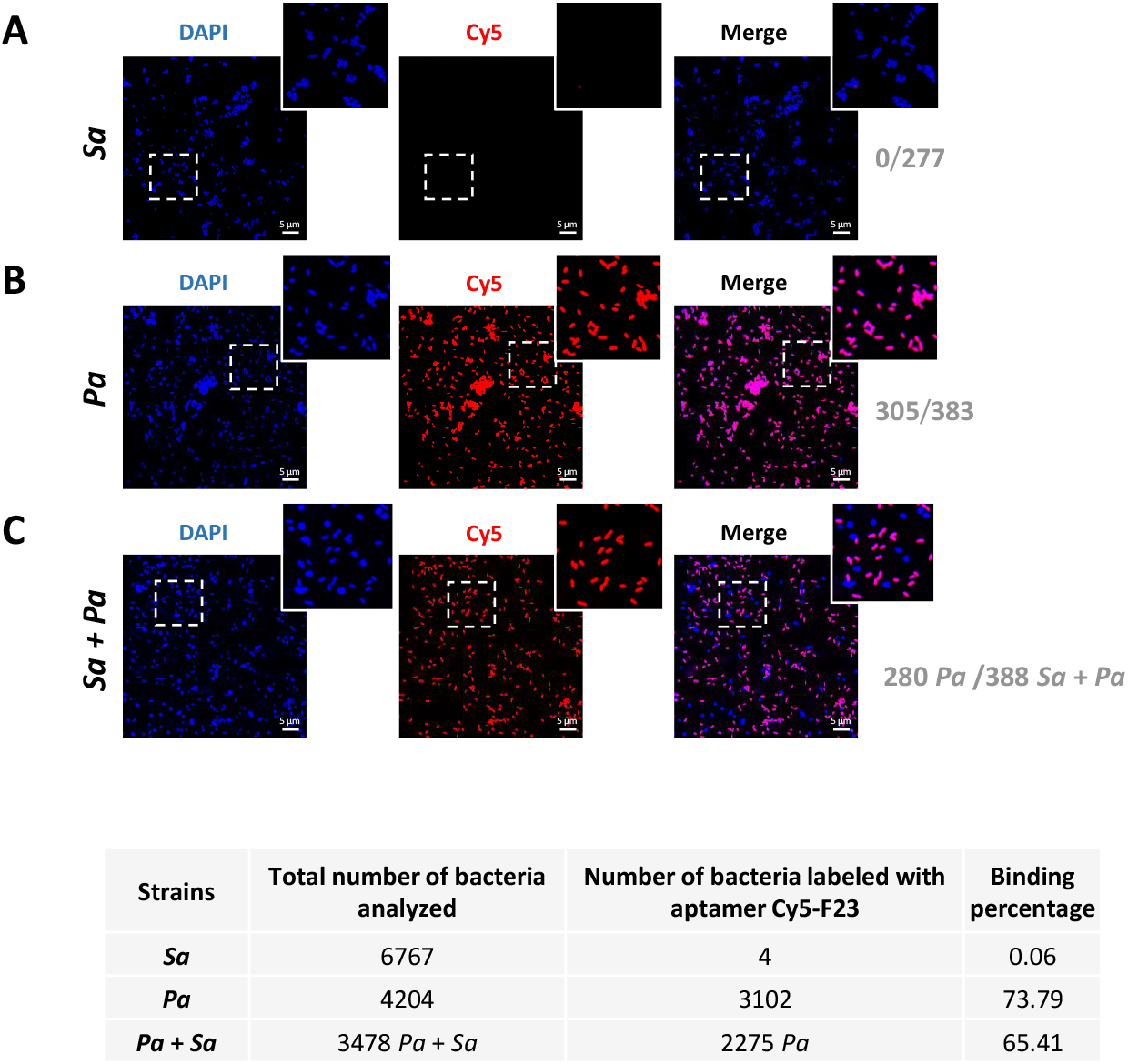
Confocal microscopy analysis of F23 aptamer on living *Pseudomonas aeruginosa* (*Pa*) and *Staphylococcus aureus* (*Sa*). **A**. Live *Sa* cells, **B**. live *Pa* cells, **C**. live *Sa + Pa* heterogeneous cocktail. Briefly, living *Sa* or/and *Pa* were incubated with 1 μM of the Cy5-F23 aptamer. Bacteria were then labelled with DAPI for visualization under microscopy (represented in blue). A positive binding of the F23 aptamer is shown in red (Cy5, λex = 644 nm). Magenta bacteria in the merged images means a positive binding of the aptamer. Note that *Pa* and *Sa* can also be differentiated by their respective shapes: bacillus and cocci, respectively. Negative binding is shown for living *Sa* alone and in the bacteria mixture. The F23 aptamer binds only to *Pa* (shown in magenta). Scale bar: 5 μm. Magnified images are from the inserts. The table presents the total number of cells analyzed, along with the corresponding aptamer-binding percentages, for isolated stains of *Sa* and *Pa*, as well as for the cocktail of *Pa*+*Sa*.

In a heterogeneous bacterial community containing both live *P. aeruginosa* and *S. aureus*, confocal images clearly demonstrate that the Cy5-F23 aptamer binds exclusively to *P. aeruginosa* (Figure 8C). This is evidenced by the distinct magenta fluorescence observed in 280 *P. aeruginosa* cells labelled by the Cy5-F23 aptamer out of a total bacteria of 388, indicating specific recognition of *P. aeruginosa* cells distinguishable by their bacillus morphology. These results underscore the high specificity of the F23 aptamer for live *P. aeruginosa*, even in the presence of other live bacterial species.

## 4. DISCUSSION

Accurately measuring bacterial populations is critical across various microbiological fields, including public health, biotechnology, food safety, water quality, pharmaceuticals, and environmental research (Lebaron et al., 1998). However, the precise identification of specific bacterial species within complex bacteria communities (heterogeneous strains) remains a considerable challenge. Nucleic-acid aptamers have not yet been extensively studied for bacterial detection. In 2011, Wang et al. (Wang et al., 2011) selected single-stranded DNA (ssDNA) aptamers targeting *Pseudomonas aeruginosa* using a whole-bacterium SELEX strategy. After 15 rounds of selection with *Pseudomonas aeruginosa* ATCC 27853 as the target, and *Stenotrophomonas maltophilia ATCC 13637* and *Acinetobacter baumannii* ATCC 17978 for negative selection steps, they identified the F23 aptamer as the most affine aptamer with a dissociation constant K_D_ of 17.27 ± 5.00 nM. The study also introduced the first application of aptamer-based fluorescence *in situ* hybridization (aptamer-FISH) for detecting ethanol-fixed bacteria, using the conjugated fluorescein isothiocyanate (FITC) F23 aptamer as the detection probe. The binding of the fluorescent F23 aptamer to *P. aeruginosa* was confirmed through hybridization against multiple *P. aeruginosa* reference strains, as well as other *Pseudomonas* species, including *Pseudomonas fluorescens, Pseudomonas putida* and *Pseudomonas stutzeri*. Additional testing against other closely related Gram-negative and Gram-positive species including *Acinetobacter baumannii, Stenotrophomonas maltophilia, Klebsiella pneumoniae, Escherichia coli* and *Staphylococcus aureus* demonstrated no detectable fluorescence. The fluorescence intensities observed for the four *P. aeruginosa* strains were similar, while no fluorescent light was detected when hybridizing with the three *Pseudomonas* strains and the five other bacterial species sus-mentioned. As the study by Wang et al. was not primarily focused on bioimaging, it included a single representative confocal fluorescence image of *P. aeruginosa* labeled with the F23 aptamer. Subsequent studies have utilized the F23 aptamer for various diagnostic and therapeutic applications, as detailed in a recent comprehensive review on aptamers targeting *P. aeruginosa* (Gutiérrez-Santana and Coria-Jiménez, 2024), but none of those are related to quantitative bioimaging.

In our study, we used the cyanine 5-conjugated F23 aptamer to detect and quantify different bacterial species through confocal imaging. The approach employed for detecting both fixed and live bacteria strains is illustrated in Figure 1. This innovative method is the first of its kind, enabling precise quantification of fluorescent aptamer binding to multiple bacteria strains. The process was supported by a quantitative analysis framework using an open-source macro, which is freely available. This macro standardizes bacterial detection in bioimaging studies involving fluorescent aptamers. Its versatility and adaptability make it a powerful tool for quantifying bacterial binding in studies utilizing other fluorescent ligands than aptamers, such as antibodies or peptides.

Our results encompass a total of 181,985 analyzed bacteria, with an average of 409 bacteria per analyzed image and a maximum of 2,492 bacteria in a single image. Zooming in on the images demonstrates that the aptamer provides a remarkably clear signal-to-noise ratio. The data show that the F23 aptamer binds with outstanding accuracy to both laboratory and clinical strains of *P. aeruginosa* whether fixed or live. A striking example is illustrated in Figure 5B, where the F23 aptamer successfully labels 1122 out of 1123 bacteria in a single image, highlighting the high sensitivity of this quantitative detection method. We also demonstrated that the F23 aptamer does not detect the Gram-positive *S. aureus, S. haemolyticus, S. epidermidis*, and *C. striatum*, as well as the Gram-negative *K. pneumonia* and *E. coli*, with detection rates below 3%. Interestingly, F23 provides thus an interesting tool to differentiate *P. aeruginosa* from *S. aureus* (one major pathogen affecting cystic fibrosis patients), and from different strains of the skin microbiota which are important in the case of infections of patients suffering from severe burns.

However, this quantitative method also revealed partial binding of the F23 aptamer to the Gram-positive *B. subtilis* (27.43%) and to the Gram-negative *S. maltophilia* (31.27%) and to a lesser extent *B. multivorans* (5.94%). The confocal images in Figure 5C show that the detection is specifically targeted at certain bacteria among others, rather than portions of the image, highlighting the consistency and precision of the technique. The binding of the Cy5-F23 aptamer to *S. maltophilia* was unexpected. The F23 aptamer was identified through the SELEX (systematic evolution of ligands by exponential enrichment) process (Ellington and Szostak, 1990). During SELEX, interfering sequences in the nucleic-acid library are removed through negative selection (also known as counter selection), while specific sequences are enriched through positive selection on the target of interest. In the study by Wang et al (Wang et al., 2011), the nucleic acid library was incubated with the target *P. aeruginosa*, and counter-selection steps were realized to avoid cross-binding with the non-target bacteria, *S. maltophilia* ATCC 13637 and *A. baumannii* ATCC 17978 during the 12th and 14th SELEX cycles. However, the strains referred to by Wang et al. (Wang et al., 2011) and in our study are different, which suggests that aptamer binding may differ between strains, even within the same species. This could also be attributed to incomplete elimination of cross-reactive sequences or similarities in surface molecular epitopes among *S. maltophilia*, and *P. aeruginosa*. This could also be attributed to the higher aptamer concentration used in our protocol (1 μM compared to 400 nM in Wang et al.) and our optimized approach, which uses PFA for fixed cells but avoids the use of alcohol. PFA fixation might indeed have a low influence on aptamer binding as detection of *P. aeruginosa* is slightly higher on PFA-fixed cells compared to live cells.

Our results also indicate that the F23 aptamer binding is not associated with bacterial classification as Gram-positive or Gram-negative, nor with bacterial morphology (round or rod-shape). Further research is needed to identify the molecular target of the F23 aptamer and to elucidate the observed and quantified differences in binding. It is also possible that some of the tested bacterial populations are not homogeneous, and that the F23 aptamer can distinguish subtle variations between strains.

Diagnostic aptamers could hold significant potential for visualizing and quantifying contaminating bacteria within ostensibly homogeneous populations or detecting pathogenic bacteria in various contexts, such as food/beverage safety, environment care, human health. Additionally, aptamers can serve as valuable tools for molecular imaging and therapeutic monitoring in bacterial infections, and even therapeutic applications (Ye et al., 2024). Despite their promise, aptamers still face notable challenges that must be addressed to enhance their performance in bacterial detection, with stability and selectivity being key limitations. First, the F23 aptamer’s stability is evidenced by its resistance to degradation (Figure 2), which is essential for preserving its binding ability and efficacy. Second, regarding selectivity, this study, along with others in different contexts such as the detection of cell-surface biomarkers (Kelly et al., 2021) or low molecular weight compounds (Bottari et al., 2020) highlights the need for further improvement to enhance aptamer performance. This could be achieved through optimization of the selection process. To address cross-reactivity and further refine the selection of aptamers, more stringent counter-selection steps have to be considered during the SELEX process, with a clear focus on aptamer ultimate intended application and usage conditions (e.g., medium, pH, temperature). This could involve using higher concentrations of non-targeted molecules or incorporating a broader range of bacterial strains and species during counter-selection. Additionally, varying environmental conditions, such as pH levels and ionic strengths, could help eliminate less selective aptamers. Structural comparisons and epitope mapping can further aid in identifying unique surface epitopes, thereby reducing the likelihood of cross-binding to non-specific strains. Targeting smaller and distinct epitopes, and accounting for glycosylation differences between bacterial strains, can help improve aptamer specificity (Kohlberger and Gadermaier, 2022; Li et al., 2021; Zhuo et al., 2017). The integration of next-generation sequencing (NGS) into the SELEX process has become a routinely practice for analyzing aptamer pools post-selection (Kolm et al., 2020). NGS allows for the refinement of aptamer pools, favoring sequences with high specificity and reduced cross reactivity. By adopting these strategies, the selectivity of aptamers can be significantly enhanced, thereby improving their utility for precise therapeutic and diagnostic applications.

## CONCLUSION

Unlike existing FISH probes, which often require costly or complex steps such as peptide nucleic acid (PNA) probes or antibodies, aptamer-FISH combines both cost-effectiveness and simplicity. Our novel approach, which integrates aptamer-FISH with quantitative bioimaging analysis, improves bacterial detection and identification, making it particularly valuable for future diagnostic applications. Since it does not require permeabilization steps, this aptamer-based method is simpler and more convenient than traditional DNA-based FISH techniques. Additionnaly, it provides a cost-effective alternative compared to PCR-based assays, which rely on target genes, primer sequences, DNA extraction procedures, and detection methods (Yamamoto, 2002). More importantly, it offers additional insights, such as cell counts, morphology and the ability to differentiate heterogeneous populations *in situ*. This quantitative aptamer-FISH method represents a promising alternative for the efficient detection of bacteria in both clinical and environmental settings. Stability remains a critical factor, and the demonstrated robustness of aptamers in complex media further highlights their potential for extended applications. Beyond diagnostics, selective aptamers are promising ligands for innovative therapeutic strategies, including targeted antimicrobial therapies and precision drug delivery systems to combat *P. aeruginosa* infections. Furthermore, our versatile aptamer-based quantitative detection platform could easily be adapted for other clinically relevant bacterial pathogens, and to other fluorescent molecular tools, broadening its applicability. By integrating this approach with advanced biosensing systems or microfluidic devices, its performance in point-of-care settings could be further optimized, paving the way for highly efficient and versatile diagnostic solutions.

## ABBREVIATIONS

BSA: bovine serum albumin
Cy5: cyanine 5
DAPI: 4’,6-diamidino-2-phenylindole 2HC1
DPBS: Dulbecco’s phosphate-buffered saline
FISH: fluorescence in situ hybridization
GFP: green fluorescent protein
MH: Mueller Hinton II medium
OD: optical density
PFA: paraformaldehyde
RT: room temperature
tRNA: transfer RNA

## ACKNOWLEDGMENT

We thank A. Doleans Jordheim, B. Tümmler, and A. Grillon for the gift of bacterial isolates. We gratefully acknowledge the Imaging Center PIQ-QuESt (https://piq.unistra.fr/), member of the national infrastructure France Bioimaging, supported by the French National Research Agency ANR-10-INSB-04.

## AUTHOR CONTRIBUTIONS

Conceptualization, P.F. and L.C.; Methodology, C.M., R.V., B.A., H.AB., M.O., P.F. and L.C.; Formal analysis, C.M., R.V., P.F. and L.C.; Investigation, P.F. and L.C.; Writing—original draft preparation, C.M., R.V., P.F. and L.C.; Writing, C.M, and L.C.; Project administration, P.F, and L.C.; Funding acquisition, P.F, and L.C. All authors have read and agreed to this version of the manuscript.

## DATA AVAILABILITY STATEMENT

Data supporting reported results will be provided by corresponding author upon request.

