## Supplemental information for "Bioimaging with fluorescent nucleic-acid aptamers for the specific detection and quantification of *Pseudomonas aeruginosa* alone and in heterogeneous bacterial populations"

### Supplementary Data 1: Tutorials

#### Installation instruction

##### 1. Install ImageJ or FIJI

Way 1 : ImageJ
- Download and install ImageJ https://imagej.net/ij/download.html
- Update ImageJ to version 1.54g or higher from the menu *Help/Update imageJ …*

Way 2 : FIJI
- Download and install FIJI  https://imagej.net/software/fiji/downloads
- Update ImageJ to version 1.54g or higher from the menu *Help/Update imageJ …*

##### 2. Install macros

Way 1
- Paste macros (*.ijm files) in the imageJ *plugins* subfolder
- Restart ImageJ to complete the process.
- Macros are now available in the Plugins menu

Way 2
- Paste macros (*.ijm files) in the imageJ *plugins* subfolder
- Navigate to Plugins -> Install -> Path to File. Restart ImageJ to complete the process.
- Macros are now available in the Plugins menu

Way 3
- Aternatively, drag and drop macro in the imageJ bar to open the macro editor
- you can run macros by clicking on the *Run* button

#### Acquisition

- Avoid saturation
- Avoid crosstalk between DAPI and Cyto9, it cause false pseudomonas detection
- If microscope do sequential acquistion, bacterias can move between acquisition of different channels. It cause a drift who can distort classification analysis (picture below)

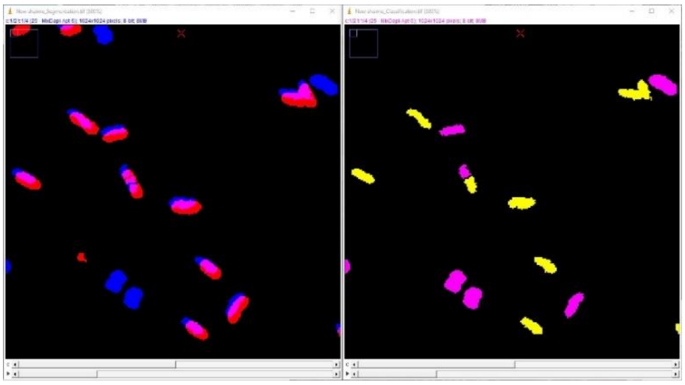

Possible solutions are :

- Use simultaneous detection of both channels
- Reduce the threshold of overlap to classify the aptamer positive cells (see in *Parameters Customisation)*

#### Parameters customisation

Segmentation parameters are set for 1024*1024 pixels image acquired with LSM confocal 63x.
Editable settings can be modified in the script : open the macro in any text editor and change variable in the *"+++ parameters +++"* paragraph, instruction are noted as comment in the script.

```
//+++++++++++++PARAMETERS ++++++++++++++++++++++++++++++++++++++++++++++++++++++++++++++++++++++++++++++++++++++++++++++++++++++++++++
    TabExt =" .tsv"     ;  // results table file format : ".xls", ".tsv", ".csv" or ".txt"

    CanDapi = 1         ;  // DAPI - Staphylococcus channel #
    CanGFP  = 2         ;  // Syto9 - Pseudomonas channel #
    CanApta = 3         ;  // Cy5 - Aptamer channel #

    BactoMinSize = 25           ;  // Bacteria minimal size in pixels
    BactoMaxSize = 300          ;  // Bacteriaa maximal size in pixels
    ClassificationSeuil = 0.6   ;  // %age of overlap with aptamer signal in a segmented bacteria to be considered as postive to aptamer labelling (between 0 and 1)

    Background = 200        ;  // background level : this value is subtracted on DAPI and Syto9 image before analysis
    Seg2Gauss = 0.5         ;  // Gaussian prefilter size
    Seg2PromIndex = 500     ;  // Prominence of maxima detection for segmentation. The lower it is the more it detect objects

    TestMode = false        ;  // developper option : switch off batchmode and keep intermediate images
//+++++++++++++++++++++++++++++++++++++++++++++++++++++++++++++++++++++++++++++++++++++++++++++++++++++++++++++++++++++++++++++++++++++
```

#### Input files

- Macros run with tif file, if images are in a microscope manufacturer format, they must be converted.
We recommand the bioformat importer tool http://www.openmicroscopy.org/bio-formats/
- Batch macro detect automatically tif files with the following extensions : *.tif , *.Tif and *.TIF

#### Output files

- Results are automatically saved in a new subfolder
- Measurement tabs are saved in *.tsv format
- Segmented images are saved in *.tif files
- Region of interest are saved in *.roi format and stacked in *zip files

### I. Aptamers Pseudo Specificity test ROI

This first classification macro was developed to test the selectivity of aptamer for Pseudomonas bacteria.

<u>Input :</u>
3 channels image :

- channel 1 : DAPI - Staphylococcus
- channel 2 : Cyto-9 - Pseudomonas
- channel 3 : CY5 - Aptamer

<u>Process :</u>

1) All channels are segmented
2) Colocalization (surface overlap) is measured between channels 1 and and 3 between channels 2 and 3
3) Cells are classified

<u>Ouput :</u>
3 channels image for segmentation (binaries) :

- channel 1 (Blue) : DAPI - Staphylococus
- channel 2 (Green) : Cyto-9 - Pseudomonas
- channel 3 (Red) : CY5 - Aptamer

4 channels image for classified bacterias (binaries) :

- channel 1 (Blue) : Staphylococus untagged with aptamer
- channel 2 (Magenta) : Staphylococus tagged with aptamer
- channel 3 (Green) : Pseudomonas tagged with aptamer
- channel 4 (Red) : Pseudomonas untagged with aptamer

Region of interests for each staphylococcus
Magenta : Staphylococus tagged with aptamer
Yellow : Staphylococus untagged with aptamer

Region of interests for each staphylococcus
Green : Pseudomonas tagged with aptamer
Red : Pseudomonas untagged with aptamer

Result table , for each bacteria are measured :
Surface
Surface labelled with aptamer
Proportion of labelled surface
Bacteria circularity
Gobal statistics

#### II. Aptamers Pseudo Specificity test Batch

This classification macro was developed to test the selectivity of aptamer for Pseudomonas bacteria. It is identical to *Aptamers Pseudo Specificity test ROI* except it run analysis on all image in a folder and region of interest are not saved.

<u>Input :</u>

A folder containing 3 channels images in tif format :

- channel 1 : DAPI - Staphylococcus
- channel 2 : Cyto-9 - Pseudomonas
- channel 3 : CY5 - Aptamer

<u>Process :</u>

1) All channels are segmented
2) Colocalization (surface overlap) is measured between channels 1 and and 3 between channels 2 and 3
3) Cells are classified

<u>Ouput :</u>

Stack image with 3 binary channels from segmentation:

- channel 1 (Blue) : DAPI - Staphylococus
- channel 2 (Green) : Cyto-9 - Pseudomonas
- channel 3 (Red) : CY5 - Aptamer

Stack image with 4 binary channels of classified bacterias:

- channel 1 (Blue) : Staphylococus untagged with aptamer
- channel 1 (Magenta) : Staphylococus tagged with aptamer
- channel 2 (Green) : Pseudomonas tagged with aptamer
- channel 3 (Red) : Pseudomonas untagged with aptamer

Result table for each image (*.tsv) with classification statistics

Result table (*.tsv) with gobal classification statistics

#### III. Aptamers Pseudo Classifier Batch

This macro identify Pseudomonas in a bacteria culture mix, where all cells are tagged with DAPI (or any another label). It runs on all tif files of a selected folder.

Input :

A folder containing 2 channels images in tif format :

- channel 1 : DAPI - Staphylococcus
- channel 2 : CY5 - Aptamer

Process :

1) All channels are segmented
2) Colocalization (surface overlap) is measured between channels 1 and and 2
3) Cells are classified

Ouput :

Stack image with 2 binary channels from segmentation :

- channel 1 (Blue) : All bacterias
- channel 2 (Red) :  Pseudomonas

Stack image with 2 binary channels of classified bacterias:

- channel 1 (Magenta ) : Bacterias untagged with aptamer
- channel 2 (Yellow) : Bacterias tagged with aptamer

Result table for each image (*.tsv) with classification statistics

Result table (*.tsv) with gobal classification statistics

#### Supplementary Data 2: Macros

Macros developed for this study can be found at and downloaded from :
https://seafile.unistra.fr/d/aac2e85b5359476e8644/

When this manuscript will be accepted for publication, these macros will be available in open access from https://git.unistra.fr
